# Hmgcr promotes a long-range signal to attract germ cells which is aided by Wunens but independent of *hh*

**DOI:** 10.1101/333575

**Authors:** Kim Kenwrick, Amrita Mukherjee, Andrew Renault

## Abstract

**Summary Statement:** Migrating *Drosophila* germ cells are attracted by a long range Hmgcr mediated signal which is aided and acts simultaneously with Wunens suggesting that these pathways converge on a single chemoattractant.

**Abstract:** In a developing embryo, many cell types migrate from their point of specification to their final position. This usually involves highly stereotyped routes which are determined through deployment of cell surface or secreted guidance molecules. Whilst genetic techniques have been successful in identifying these molecules, the distances over which such signals operate in their native context can be difficult to determine. Here we have quantified the range of an attractive signal for the migration of *Drosophila* germ cells. Their migration is guided by an attractive signal generated by the expression of genes in the 3-hydroxy-3-methyl-glutaryl-coenzyme A reductase (Hmgcr) pathway, and by a repulsive signal generated by the expression of Wunens. We demonstrate that the attractive signal downstream of Hmgcr operates over a long range and is sufficient to reach germ cells for the entirety of their migration. Furthermore, Hmgcr-mediated attraction and Wunen-mediated repulsion can operate simultaneously ruling out a model in which these pathways operate consecutively. Indeed, we show that Hmgcr-mediated attraction is boosted by Wunens suggesting the action of these two pathways is linked. Lastly, several papers have pointed to the secreted molecule Hedgehog (Hh) as being the germ cell attractant, whose secretion is increased by *hmgcr*. In this paper, we provide evidence that Hh is not downstream of *hmgcr* in germ cell migration.

## Introduction

Cells are often on the move. Microorganisms migrate to find nutrients or a suitable host. Cells in developing embryos can be swept around via large morphogenetic movements, or move more subtly either individually or as small collectives of cells pushing through and between tissues. Cells find their way by detecting the presence and/or concentration of secreted or cell surface molecules which act as either chemoattractants or chemorepellants. Chemoattractants may be secreted by destination tissues and also by cells along the migratory route that act as intermediate targets. Localised destruction or uptake of chemoattractants are often important for shaping the gradients as well as encouraging cells to leave the intermediate staging points (Yu et al., 2009; Boldajipour et al., 2008). Cells may also use multiple chemoattractants simultaneously such as in the case of border cells in the *Drosophila* ovary (Duchek and Rørth, 2001; Duchek et al., 2001).

One cell type, whose migration has been studied extensively is the primordial germ cells. These are the cells that give rise to the gametes in adults. They are formed early in development and migrate during embryogenesis to the gonad in many, but not all, model organisms (reviewed in Barton et al., 2016). Their prominence as a model for cell migration arises from their importance for continuation of the species, ease of identification by morphology, embryonic position and gene expression profile and highly stereotyped migratory routes.

The *Drosophila* primordial germ cells are initially moved by gastrulation from their site of formation at the posterior pole into the posterior midgut pocket. Migration begins with the germ cells pulling away from each other and traversing the posterior midgut (Seifert and Lehmann, 2012) (Fig 1A stage 10). They then move towards the dorsal side of the midgut epithelium and enter the overlying mesoderm, partitioning bilaterally as they do so (Sano et al., 2005) (Fig 1A stage 11). In the mesoderm they associate with the somatic gonadal precursors (SGPs) (Fig 1A stage 12), at which point their migration ceases, and together they coalesce to form the embryonic gonad (Boyle and DiNardo, 1995) (Fig 1A stage 14).

**Figure 1.**
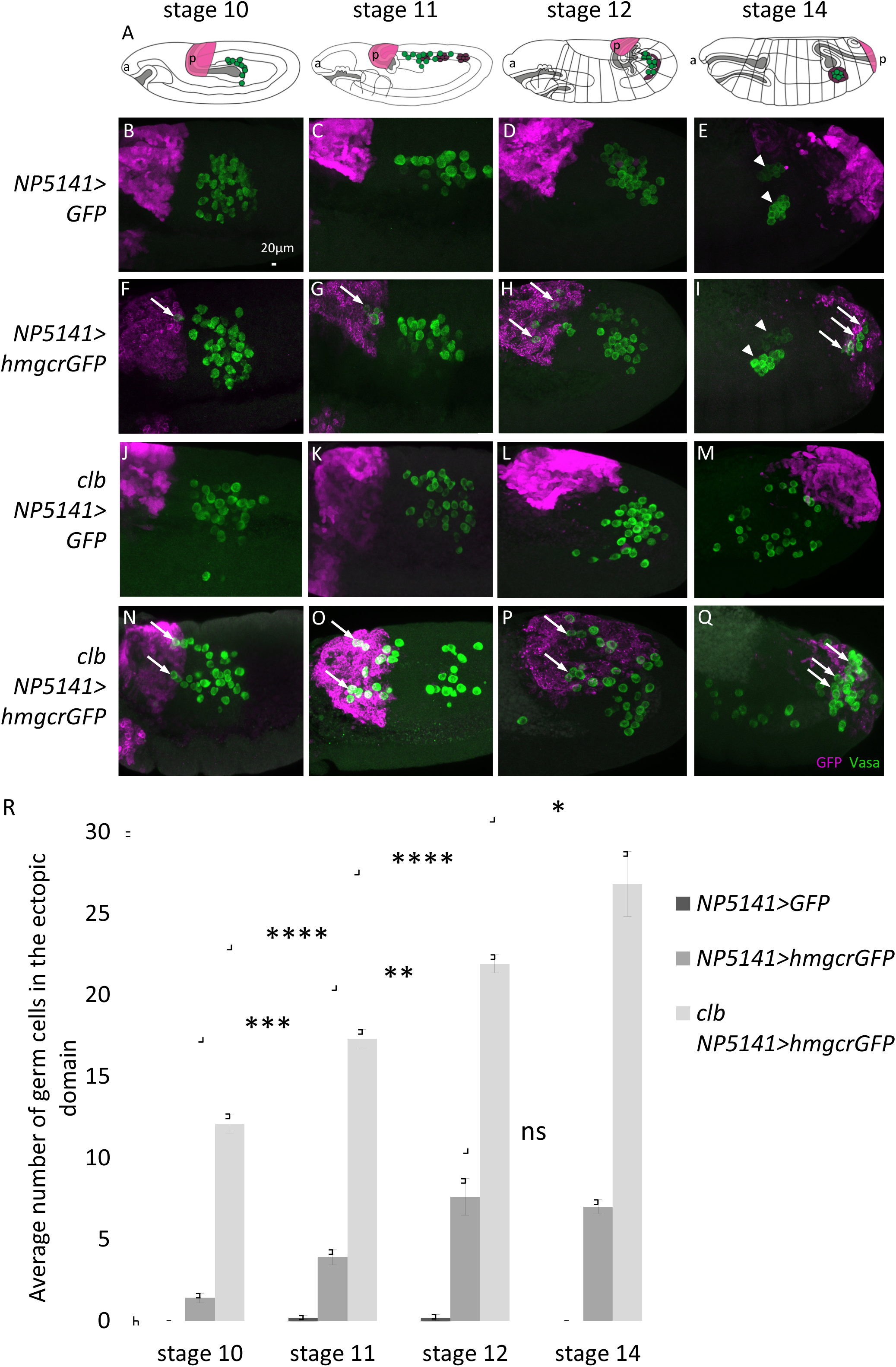
*hmgcr* expressing endogenous and ectopic domains compete to attract germ cells. (A) Cartoon of lateral views, with anterior (a) to the left, of *Drosophila* embryos at the stage indicated showing germ cells (green) and SGPs (purple) relative to the *NP5141Gal4* domain in the most posterior parasegment (magenta). Following germ band extension, a stage 10 embryo is folded over on itself so the posterior (p) lies above and slightly to the left of centre. During stage 12 the germ band retracts pulling the posterior to its final position on the right. (B-Q) Maximum intensity projections of lateral views of representative embryos, of the genotypes: (B-E) *NP5141Gal4/UASGFP,* (F-I) *NP5141Gal4/+; UAShmgcrGFP/+,* (J-M) *NP5141Gal4/UAS GFP; clb11.54/clb11.54,* (N-Q) *NP5141/+; clb11.54 UAShmgcrGFP / clb11.54* at stages 10-14, fluorescently stained with antibodies against Vasa to label the germ cells (green) and GFP to visualize the ectopic domain (magenta). Arrowheads indicate the position of the embryonic gonads and arrows indicate germ cells that have been attracted to the ectopic domain. (R) Graph showing the mean ± s.e.m number of germ cells in the ectopic domain, in embryos at stages 10-14 with n=10 embryos scored per genotype

Forward genetic screens in *Drosophila* have identified two important enzymatic pathways for germ cell migration. The first comprises enzymes of the 3-hydroxy-3-methylglutaryl coenzyme A (Hmgcr) pathway which catalyzes the conversion of acetyl groups to the isoprenoids farnesyl- and geranyl geranyl-pyrophosphate which are used for protein prenylation, as well as being precursors for other lipids such as juvenile hormone (Bellés et al., 2005). Mutations in *columbus* (*clb*), the *Drosophila* gene encoding Hmgcr, cause germ cells to scatter over the posterior of the embryo (Van Doren et al., 1998). Hmgcr is expressed broadly in the mesoderm before becoming enriched in just the mesodermally derived target tissue, the SGPs (Van Doren et al., 1998). Ectopic expression of this enzyme, in tissues such as the CNS or the ectoderm, is sufficient to attract a small number of germ cells into the tissue of ectopic expression (Van Doren et al., 1998; Ricardo and Lehmann, 2009). These data suggest that the Hmgcr pathway is involved in producing a chemoattractant that attracts the germ cells to the SGPs (Fig S1).

Some studies report that Hedgehog (Hh) is the Hmgcr-dependent germ cell attractant (Deshpande and Schedl, 2005; Deshpande et al., 2001; Deshpande et al., 2013). However Hh itself is not prenylated (Eaton, 2008) and the ability of Hh to attract germ cells has not proved reproducible (Renault et al., 2009). Therefore, the identity of the chemoattractant molecule downstream of Hmgcr remains controversial.

The second pathway comprises two enzymes, Wunen and Wunen2, hereafter collectively referred to as Wunens. The Wunens are lipid phosphate phosphatases (LPPs), integral membrane enzymes that can dephosphorylate and internalize extracellular lipid phosphates (Sigal et al., 2005). The Wunens are expressed in somatic regions that germ cells do not normally enter and in the absence of this expression, germ cells scatter over the posterior of the embryo (Starz-Gaiano et al., 2001; Zhang et al., 1997). Overexpression of Wunens blocks germ cell entry into the ectopic tissue and induces many germ cells to die (Starz-Gaiano et al., 2001). In a purely phenomenal (but not necessarily a mechanistic) sense, Wunen expression can be thought of as repelling germ cells (Fig S1). Wunens are also expressed on germ cells themselves (Hanyu-Nakamura et al., 2004; Renault et al., 2004) and this leads to germ cell-germ cell repulsion that may be responsible for their initial dispersal out of the posterior midgut (Renault et al., 2010) (Fig S1).

The prevailing idea is that Wunens act as a sink for an extracellular lipid phosphate attractant (Renault et al., 2004). Although this extracellular lipid phosphate attractant has not been identified for *Drosophila*, in the ascidian *Botryllus schlosseri*, sphingosine 1-phosphate (S1P) can direct germ cell migration. S1P is an *in vitro* substrate for LPPs (Roberts et al., 1998) raising the possibility that S1P, or a related molecule, acts as an attractant in *Drosophila*. Recent work demonstrates that the signal downstream of Wunens is likely perceived by germ cells via Tre1, a G-protein-coupled receptor (GPCR) on germ cells (LeBlanc and Lehmann, 2017).

The most recently proposed model of germ cell migration supposes that the Hmgcr and Wunen pathways work independently of each other (Barton et al., 2016). SGPs produce a prenylated germ cell attractant via action of the Hmgcr pathway. This prenylated attractant is perceived by germ cells via an unidentified receptor and acts as an attractant. Wunens create a separate gradient of extracellular S1P or a related lipid which also acts as a germ cell attractant (Barton et al., 2016). The Tre1 GPCR on germ cells is responsible for sensing the substrate of the Wunens (LeBlanc and Lehmann, 2017) leaving the identity of the germ cell receptor for the Hmgcr-dependent chemoattractant unknown.

Such a model leaves several open questions. Firstly, do the two chemoattractants operate with similar or different characteristics? Perhaps one is a long-range signal to get the germ cells initially moving in the right direction from the midgut whilst the other acts at short range to finesse the later migration to the SGPs. Secondly, how do germ cells integrate these two signals? For example, how would germ cells respond when given conflicting guidance information by these two pathways? Perhaps in this scenario one pathway is dominant over the other.

Previously we showed that Wunens expressed in somatic cells repel germ cells without the need for cell-to-cell contact over at least a distance of 33µm, implying they regulate a long-range diffusible signal (Mukherjee et al., 2013). In this paper, we have used germ cell positioning relative to ectopic *hmgcr* expression in fixed embryos, and trajectories of migration from live imaging to obtain quantitative information on the range of the Hmgcr-dependent signal. We show that, like Wunens, the Hmgcr-dependent signal also acts at long range and can attract germ cells at distances of up to 51µm. We have also used epistatic analyses to investigate the relationship between the *hmgcr* pathway, *wunens* and *hh*. Firstly, we find that *hh* does not act downstream of *hmgcr* in attracting germ cells. Secondly, we find that removal of Wunens reduces the ability of *hmgcr* to attract germ cells. We use these data to propose a new model of germ cell migration which posits a single chemoattractant whose extracellular level is modulated by both Wunens and Hmgcr.

## Materials and Methods

### Fly Stocks

The following *Drosophila* lines were described previously: *Df(2R)wunGL*, a deficiency removing *wun* and *wun2* (Zhang et al., 1996); *clb11.54*, a loss of function allele of *hmgcr* (Van Doren et al., 1998); *hhAC*, an amorphic allele resulting from a 8.6kb deletion removing the promoter and part of the coding region (Lee et al., 1992); *UAS wun2myc* (Starz-Gaiano et al., 2001); *UAS wunGFP* (Burnett and Howard, 2003), *UAS lazGFP* (Garcia-Murillas et al., 2006), expression of which does not affect germ cells (Mukherjee et al., 2013); *UAS hmgcr* (Van Doren et al., 1998); *HmgcrEY04833*, a UAS containing insertion 5’ of the *Hmgcr* gene (Bloomington Stock Centre); *p(GawB)NP5141*, a Gal4 containing insertion 5’ of the gene *ken* (Drosophila Genetic Resource Center); *y M{vas-int.Dm}ZH-2A w; PBac{y+-attP-3B}VK00033* used as a landing site for the *UAShmgcrGFP* transgene. *nanos>moeGFP* was used to label the germ cells for live imaging (Sano et al., 2005).

### Immunohistochemistry and imaging

Embryos were laid at room temperature, dechorionated in 50% bleach for 3 minutes, fixed for 20 minutes in 4% formaldehyde (37% for *in situ* hybridisation) in PBS/heptane, devitellinized using heptane/methanol, and stained using standard protocols. Primary antibodies were as follows: rabbit anti-Vasa (1:10,000) courtesy of Ruth Lehmann, rabbit anti-LacZ (Capell 1:10,000), chicken anti-GFP (Abcam ab13970, 1:1000). Secondary antibodies conjugated to Alexa Fluor 488 or 648 (Invitrogen) and Cy3 (Jackson ImmunoResearch) were used at 1:500.

For fluorescent *in situ* hybridisation a full length *hmgcr* cDNA clone in a pNB40 vector was linearised and used to make a digoxygenin labelled RNA probe by *in vitro* transcription, and hybridisation was carried out as described previously (Lécuyer et al., 2008).

Fluorescently stained embryos were either mounted in aquamount (Polysciences) or dehydrated in methanol and mounted in benzylbenzoate: benzyl alcohol (2:1). Images were acquired using an LSM 880 confocal microscope with a x20/NA 0.5 air or x40/NA 1.3 oil objective and Zeiss Zen2 acquisition software. 3D reconstructions, segmentations and distance measurements were made with Imaris (Bitplane). For distance measurements, germ cell positions were detected automatically (using the spots tool with a spot diameter of 7um) and manually edited for accuracy. The ectopic domain was segmented using the surfaces tool. The distance of the edge of each spot (using the Imaris minimum intensity statistic) to the nearest ectopic domain surface was measured using the MeasurementPro extension.

Live imaging was performed on a Zeiss Z1 light sheet microscope and cells were tracked using Imaris.

### Generation of *UAShmgcrGFP* flies

The *hmgcr* coding sequence was amplified from cDNA clone in pNB40 using the primers CACCATGAGGACGTTTGTTTCGC and GCTGATGGGCTGCAGCTGG and cloned into the pENTR/*D-TOPO* vector (Invitrogen). The sequence was verified and moved into the destination vector pUAST-*attB*-WG (a gift from Saverio Brogna, producing C-terminal GFP fusions) with the use of the Gateway reaction. This resulting expression vector pUAST-*attB*-*hmgcr*-WG was microinjected into embryos containing phiC31 integrase and an *attP* site on the 3rd chromosome.

## Results

### Ectopic *hmgcr* expression is sufficient to attract germ cells into the ectopic domain

To address the question of whether *hmgcr* produces a short- or long-range signal to germ cells we wanted to examine the distances that germ cell migrate in order to enter domains of ectopic *hmgcr* expression (hereafter termed the ectopic domain). We therefore constructed a tagged UAS *hmgcr* overexpression construct allowing us to simultaneously attract germ cells and visualize the region of misexpression. Ectopic expression of *hmgcr GFP* was as effective at disrupting germ cell migration as previously described untagged *hmgcr* constructs indicating that the Hmgcr GFP fusion protein was functional (Fig S2).

We next wanted to ascertain if *hmgcr* expression could attract germ cells into ectopic domains as was suggested previously using CNS and ectodermal Gal4 lines (Van Doren et al., 1998). We made use of a Gal4 driver line termed NP5141 that we have previously used to measure the repulsive forces exerted by the Wunens (Mukherjee et al., 2013). We found that *hmgcrGFP* expression in the NP5141 domain is sufficient to attract germ cells away from their normal migration route and to enter the ectopic domain (Fig. 1I, compared to 1E).

In order determine the time points during which the germ cells were attracted we examined the number of germ cells in the ectopic domain at different embryonic stages. Germ cells were visualized inside the ectopic domain as early as stage 10 when germ cells have just crossed the posterior midgut and are starting to enter the mesoderm (Fig. 1F compared to 1B). Between stages 10 to 11, and 11 to 12 there were significant increases in the number of germ cells located in the ectopic domain (Fig. 1G-I compared to 1C-E, quantified in 1R) indicating that germ cell attraction to the ectopic domain occurs continually rather than at a discrete timepoint. However, between stages 12 to 14 there was no significant increase in the number of germ cells in the ectopic domain (Fig.1R). It is at these stages that in wild type germ cells make contact and start coalescing with the SGPs suggesting that once germ cells have reached the SGPs they no longer can be attracted to the ectopic domain.

### Ectopic and endogenous domains of *hmgcr* compete to attract germ cells

In the previous experiment germ cells migrate either to the SGPs (which naturally express *hmgcr*) or to the ectopic *hmgcr*. In this scenario, the SGP and ectopic *hmgcr* expressing domains are likely competing with each other to attract germ cells. This is important because it may lead us to underestimate the attractive range of the *hmgcr*-mediated signal because potentially more germ cells would be attracted to the ectopic domain were it not for the SGP *hmgcr* expression.

To test this hypothesis we expressed *hmgcr* using the NP5141 driver in a *columbus (clb)* null background (*hmgcr* loss of function alleles are termed *clb* (Van Doren et al., 1998)). We found that the number of germ cells in the ectopic domain was significantly increased compared to the wild type background at all stages (Fig 1N-Q compared to 1F-I, quantified in 1Q). Furthermore, the increase in germ cell number inside the ectopic domain continued past stage 12, unlike in the wild type background (Fig. 1R). Therefore, germ cells can continue to migrate and be attracted to the ectopic domain even late into embryogenesis in the absence of the SGP expression of *hmgcr*.

We conclude firstly that ectopic *hmgcr* does indeed compete with endogenous *hmgcr* in germ cell attraction and secondly that the temporal limit of germ cell attraction in wild type embryos is due to attraction and capture by the SGPs rather than a stage-dependent shut down of the germ cell migratory program.

### Wunen expression aids *hmgcr*-mediated germ cell attraction

Given that endogenous *hmgcr* restricts the number of germ cells that can be ectopically attracted we wanted to test whether other regulators of germ cell migration also have this effect. We therefore examined germ cell attraction in the background of a deficiency that removes somatic *wun* and *wun2* (hereafter referred to as a *wun* mutant background).

Surprisingly we found a significant reduction in the number of germ cells in the ectopic *hmgcr* domain in a *wun* mutant background compared to wild type (Fig. 2B compared to 1I, quantified in 2D). This reduction is made more compelling because in *wun* mutants just expressing GFP in the ectopic domain some germ cells stray into the posterior of the embryo due to random mismigration (Fig 2A and quantified in the first column of 2D) as has previously been observed (Mukherjee et al., 2013).

**Figure 2.**
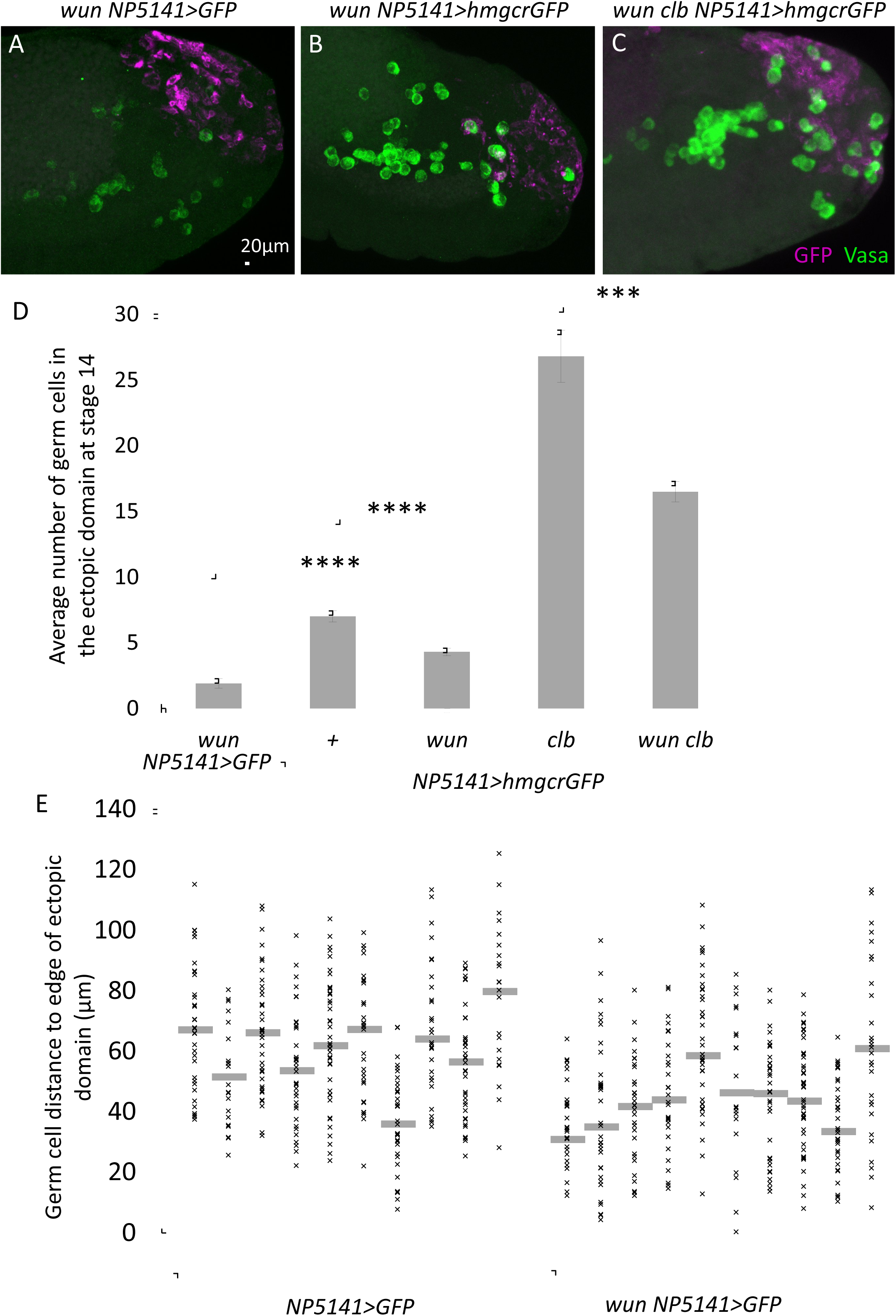
Wunen aids *hmgcr*-mediated germ cell attraction. (A-C) Maximum intensity projections of lateral views of representative stage 14 embryos, of the genotypes: (A) *Df(2R)wunGL UASlazGFP/Df(2R)wunGL NP5141Gal4,* (B) *Df(2R)wunGL/Df(2R)wunGL NP5141Gal4; UAShmgcrGFP/+,* (C) *Df(2R)wunGL/Df(2R)wunGL NP5141Gal4; clb11.54 UAShmgcrGFP/clb11.54* fluorescently stained with antibodies against Vasa to label the germ cells (green) and GFP to visualize the ectopic domain (magenta). LazGFP was used as a control protein for labelling the ectopic domain. (D) Graph showing the mean ± s.e.m number of germ cells in the ectopic domain in stage 14 embryos. n=10 embryos scored per genotype. (E) Graph showing the distances from each germ cell to the nearest surface of the ectopic domain in stage 10 embryos. Each column represents one embryo. Grey bars indicate the median distance.

To determine if Wunens also aid ectopic attraction in a *clb* mutant background we examined germ cell attraction in a triple mutant background (loss of function for *wun, wun2* and *clb*). There was also a significant reduction in the number of germ cells in the ectopic domain in the triple mutant compared to the *clb* mutant alone (Fig. 2C compare to 1Q, quantified in 2D). We conclude that somatic *wun* expression actually aids *hmgcr*-mediated attraction rather than hinders it.

We reasoned that the beneficial effects of *wun* expression might be because in *wun* mutants the germ cells are already mismigrating as they cross the posterior midgut at stage 10 and are potentially further away from the ectopic domain compared to wild type. They might therefore be less able to be attracted simply because of this increased distance. To test this hypothesis, we measured the distance of every germ cell to the nearest surface of the ectopic domain which had been computationally segmented. We found that germ cells, as they crossed the posterior midgut at stage 10, were actually closer to the ectopic domain in *wun* mutant embryos compared to wild type (Fig 2E). We conclude that loss of Wunens does not increase the distance of germ cells to the ectopic domain. Therefore, distance alone is not sufficient to explain why Wunens are beneficial for *hmgcr*-mediated attraction.

### The *hmgcr*-mediated signal is long range

We next wanted to make a quantitative assessment of the effective range over which the *hmgcr*-mediated signal attracts germ cells. Our rationale was to determine how far germ cells are from the ectopic domain when labelled with just GFP because it is at those distances that some germ cells would be attracted when it expresses *hmgcrGFP*.

We focused on stage 10 embryos when the germ cells are first attracted to the ectopic domain. Firstly, we asked how many germ cells are in the ectopic domain in experimental embryos in which the ectopic domain expresses *hmgcrGFP*. In a wild type background this was on average 2 germ cells (Fig 3C). Secondly, we took control embryos in which the ectopic domain expresses GFP and measured the distance of every germ cell to the nearest surface of the ectopic domain which had been computationally segmented (Fig 3A,B). We then recorded the distance of the 2nd closest germ cell based on the assumption that it is the two closest germ cells that would have been attracted to ectopic domain if it would have expressed *hmgcrGFP*. The distances of these 2nd closest germ cells were then averaged for the 10 embryos examined (Fig 3D).

**Figure 3.**
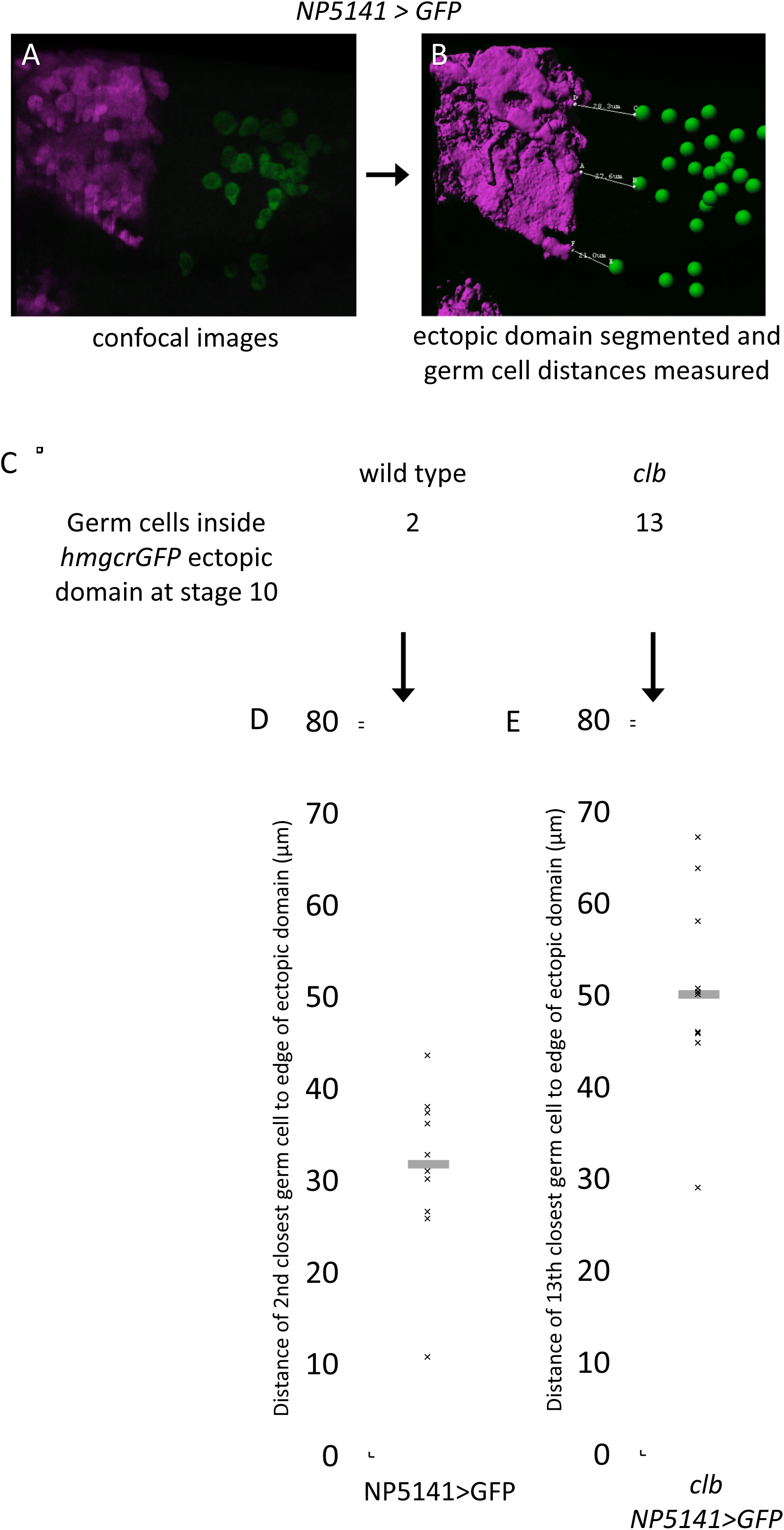
Hmgcr mediates a long-range signal. (A) Maximum intensity projection of a lateral view of a representative *NP5141Gal4/UASGFP* stage 10 embryo fluorescently stained with antibodies against Vasa to label the germ cells (green) and GFP to visualize the ectopic domain (magenta). (B) Representation of embryo in (A) after segmentation of the ectopic domain (magenta surface), assignment of germ cell positions (green spheres) and automatic measurement of distances between germ cell and ectopic domain boundaries (white lines). (C) Table showing average number of germ cells inside the *hmgcr* expressing ectopic domain at stage 10 (taken from Fig 1R) in a wild type and *clb* mutant background. (D) Graph showing the distance of the 2nd closest germ cell from a GFP expressing ectopic domain in a wild type background. The assumption is that it is these closest 2 germ cells that would be attracted to a *hmgcr* expressing ectopic domain in a wild type background and thus provides an estimate of the attraction range of ectopic *hmgcr* when in competition with wild type expression. (E) Graph showing the distance of the 13th closest germ cell from a GFP expressing ectopic domain in a *clb* mutant background. The assumption again is that it is these closest 13 germ cells that would be attracted to a *hmgcr* expressing ectopic domain in a *clb* mutant background and thus provides an estimate of the attraction range of *hmgcr* without competition from wild type expression. Grey bars indicate the median distance. Number of embryos scored = 10.

In a wild type background the second closest germ cell was on average 31µm from the ectopic domain, leading us to conclude that the *hmgcr*-mediated signal is able to attract germ cells over at least this distance.

We next considered whether we might be underestimating the effective range of the *hmgcr*-mediated signal, because germ cells in wild type embryos are still subject to competition from endogenous Hmgcr in the SGPs (Fig 1). Therefore, we applied the same methodology as described above to *clb* mutant embryos. In this case, there were on average thirteen germ cells in the ectopic domain expressing *hmgcrGFP* in *clb* mutant embryos (Fig 3C). In a *clb* mutant embryo with the ectopic domain expressing GFP alone, the thirteenth closest germ cell was on average 51µm from this domain (Fig 3E). This led us to conclude that the *hmgcr*-mediated signal is able to attract germ cells over at least 51µm. We conclude that Hmgcr acts to produce a long-range signal, and by implication cell-contact independent, in *Drosophila* embryos which attracts germ cells.

### Germ cells migrate to the ectopic domain in living embryos

So far, we have estimated the range of the *hmgcr*-mediated signal by analysing germ cells in fixed embryos. To see if we could observe germ cells being ectopically attracted over such distances in living embryos we used light sheet microscopy, which enables us to image all of the germ cells migrating in an embryo throughout embryogenesis. We visualized migrating germ cells using a *nanos>moeGFP* construct (Sano et al., 2005).

In a control embryo, in which the amnioserosa was labelled with GFP to visualise the germ band movements of the embryo, the germ cells moved from the posterior midgut pocket to the gonad over a period of approximately 6 hours. The path of migration and the lack of noticeable germ cell death indicates that the embryos were not adversely affected under the imaging conditions used (Fig 4A).

**Figure 4.**
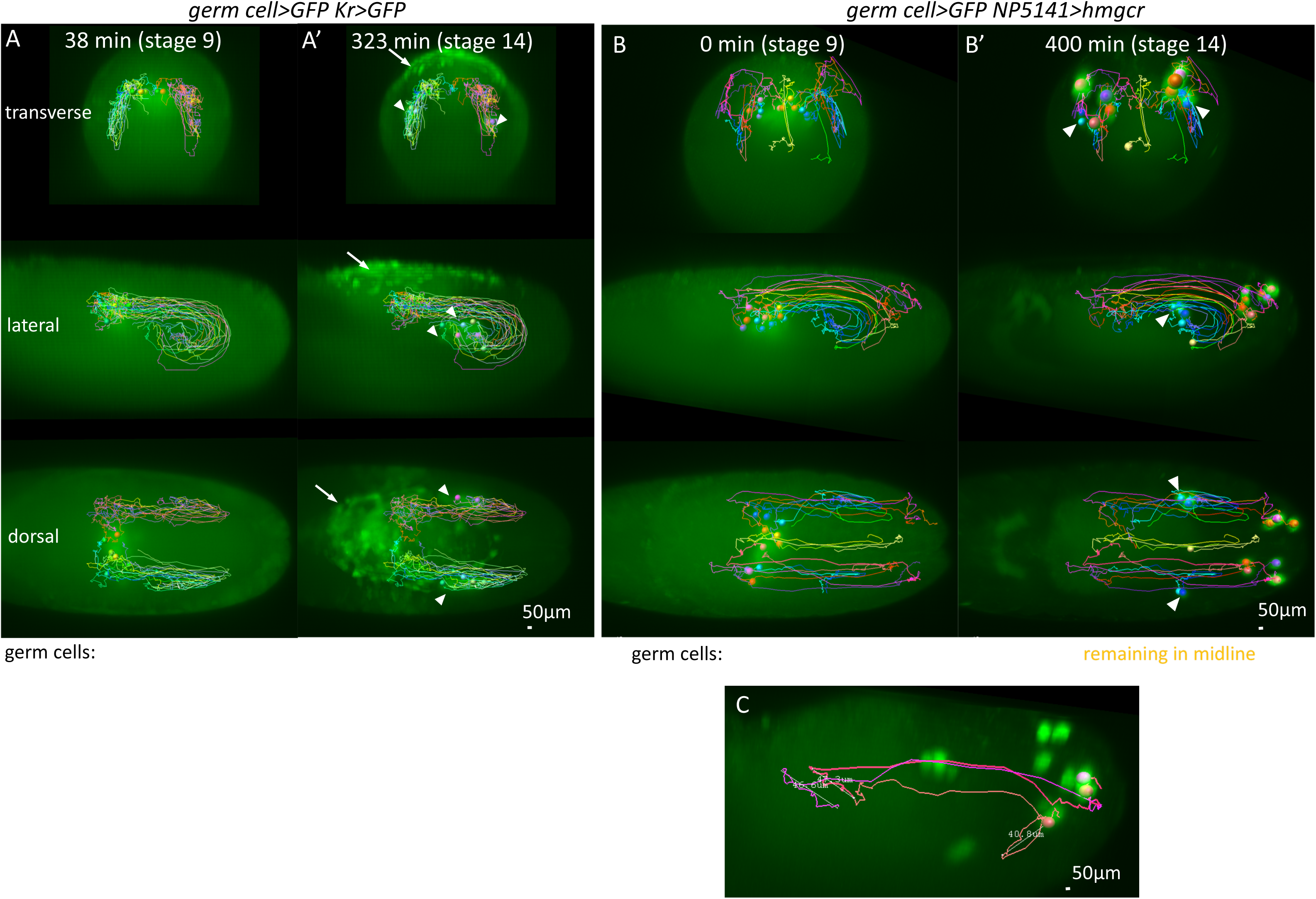
Live imaging of ectopic Hmgcr attracting germ cells. (A, B) Transverse (upper) lateral (middle) dorsal (lower) views from 3D reconstructions of movies of embryos with the genotypes: (A) *nos>moeGFP;; Pr Dr/TM3 KrGal4 UASGFP*, (B) *nos>moeGFP; NP5141Gal4/+; UAShmgcr/+.* Germ cells were tracked and coloured according to their final position. (C) Lateral view taken from embryo in (B) showing germ cell tracks for three germ cells that ended up in the ectopic domain but entered it at different stages. Linear distance between germ cell position when it first began moving into the ectopic domain and when it stopped migrating as it entered the ectopic domain are indicated.

In an experimental embryo, in which the ectopic domain expressed untagged *hmgcr* to avoid interference with the germ cell labelling, we observed germ cells migrating to the ectopic domain. We tracked the majority of the germ cells and colour coded them according to whether they migrated to the gonad as in wild type (Fig 4B, blue/cyan tracks), or to the ectopic domain (Fig 4B, pink/purple tracks) or that remained at the midline (Fig 4B, yellow tracks). We saw that germ cells started entering the ectopic domain from late stage 10 and continued to enter until late stage 12 (Fig 4B, C). Germ cells once associated with the gonad at stage 13 remained there and did not exit and migrate to the ectopic domain. These observations are in agreement with those from the fixed embryo analysis. We were also able to ascertain that germ cells once in the ectopic domain remained there and stopped migration indicating that high levels of *hmgcr* even in non-SGP somatic cells are sufficient to stop the migratory program of the germ cells (Fig 4B).

We then focused on the portions of migratory movements of germ cells entering the ectopic domain in which the germ cells broke away from the normal migratory path. We measured the distance over which this abnormal migration took place (Fig 4C). At stage 11 we observed two germ cells each migrating for approximately 47µm to enter the ectopic domain and at stage 12 we observed one germ cell migrating 41µm to enter the ectopic domain (Fig 4C). These distances are in strong agreement with our estimates of a range of 51µm from fixed embryos (Fig 3D) and confirm that *hmgcr* is mediating a long-range signal.

### Germ cells are within range of the *hmgcr*-mediated signal throughout their migratory journey

To see how our estimate for the range of the *hmgcr*-mediated signal compares to the distance of germ cells to *hmgcr* expressing SGPs in wild type embryos we measured such distances in stage 10 and 11 embryos (Fig 5A-C). We found that germ cells were located between 5 and 58µm from their closest SGP at stage 10 and ranged from 0 to 30 µm at stage 11. Therefore, for stages 10 and 11, 98% and 100% of germ cells respectively would be within our estimate of 51µm for the range of the *hmgcr*-mediated signal. We conclude that germ cells are potentially under the influence of the *hmgcr*-mediated signal for their entire migratory journey once they have left the midgut.

**Figure 5.**
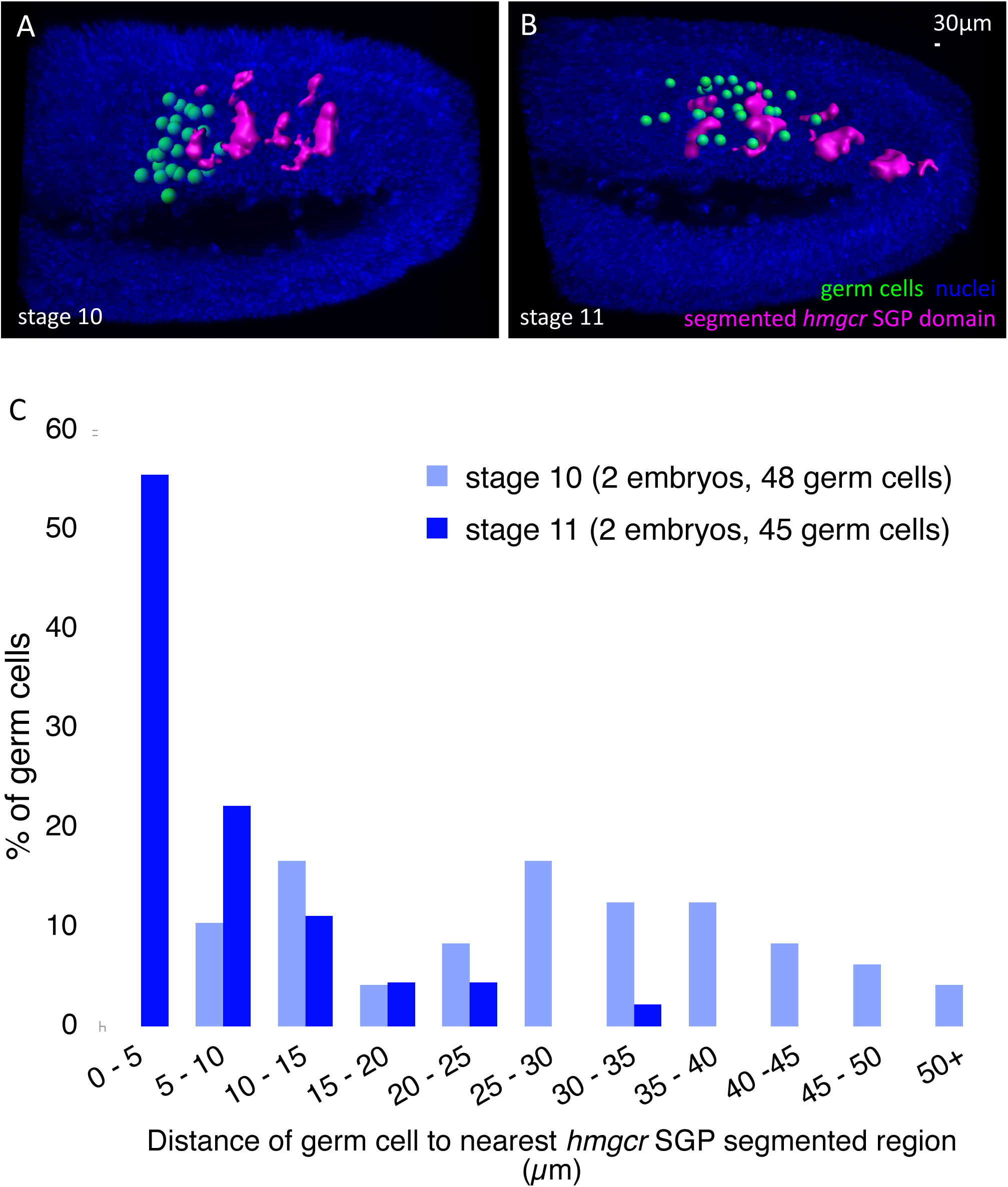
The majority of germ cells are within the range of the *hmgcr*-mediated signal. (A, B) Lateral views of 3D reconstructions of stage 10 (A) and stage 11 (B) wild type embryos that had been fluorescently stained with antibody against Vasa to label the germ cells, a RNA probe for *hmgcr* and DAPI to label nuclei (blue). Germ cells positions were manually scored (green spheres) and *hmgcr* expressing SGP clusters were computationally segmented (magenta). Other *hmgcr* expressing domains were not segmented. (C) Graph showing frequency of germ cells being located at distances indicated from their nearest *hmgcr* expressing SGP cluster at stages 10 and 11.

### *hmgcr* and *wun* operate simultaneously

We next wanted to know which of the two pathways, Hmgcr or Wunen, is dominant. To do this we gave germ cells conflicting guidance cues by simultaneously attracting them to the ectopic domain using *hmgcr* expression and repelling them by co-expressing *wun*. When *wun* is expressed using the NP5141Gal4 driver there is no effect on overall germ cell migration (Mukherjee et al., 2013 and Fig 6A-D). When *wun* and *hmgcr* are co-expressed germ cells are still attracted towards the ectopic domain similar to expression of *hmgcr* alone (Fig 6E-H). However, despite some germ cells arriving at the ectopic domain as early as stage 11 (Fig 6F), germ cells are not subsequently found within the ectopic domain but instead remain at its boundary (Fig 6G,H).

**Figure 6.**
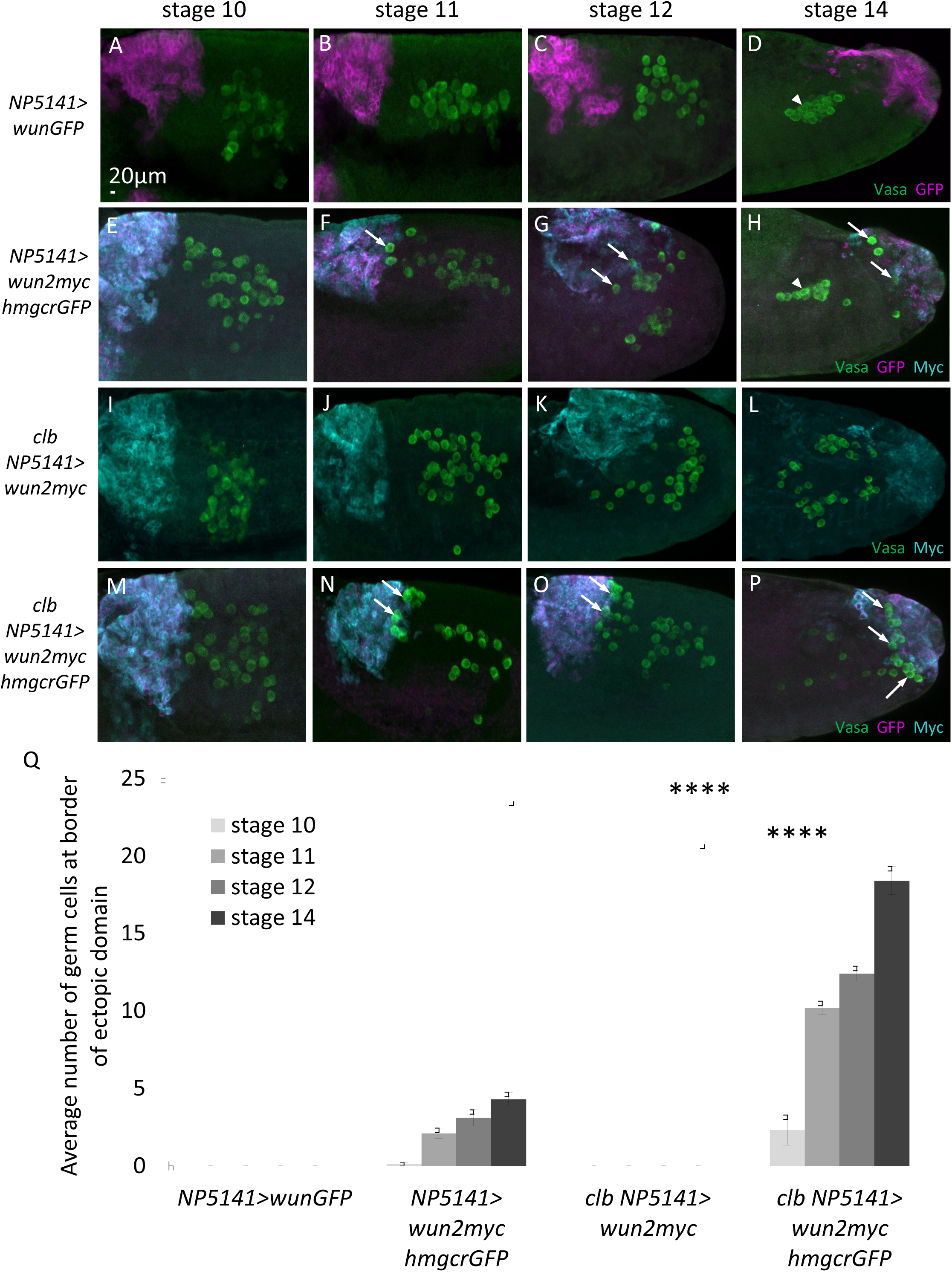
The Hmgcr and Wunen pathways can operate simultaneously. (A-P) Maximum intensity projections of lateral views of representative stage 10-14 embryos, of the genotypes: (A-D) *NP5141Gal4/UASwunGFP,* (E-H) *NP5141Gal4/UASwun2myc; UAShmgcrGFP/+,* (I-L) *NP5141/UASwun2myc; clb11.54/clb11.54*, (M-P) *NP5141/UASwun2myc; UAShmgcrGFP clb11.54/clb11.54* fluorescently stained with antibodies against Vasa to label the germ cells (green), GFP and Myc to visualise ectopic expression (magenta and cyan respectively). Arrowheads indicate the position of the embryonic gonads and arrows indicate germ cells that have been attracted to the ectopic domain. (Q) Graph showing the mean ± s.e.m number of germ cells in the ectopic domain in stage 10-14 embryos. n=10 embryos scored per genotype.

This positioning of the germ cells could result from attraction to the ectopic domain by *hmgcr* and then *wunen* activity either repelling germ cells from entering the domain or killing those germ cells that do enter, as happens for example when Wunens are ectopically expressed in the mesoderm (Starz-Gaiano et al., 2001). To distinguish between these two hypotheses, we tested what would happen if larger numbers of germ cells could be attracted to the ectopic domain. We therefore performed the same experiment in a *clb* mutant background in which competition for attraction by the SGPs is eliminated. In this scenario, the number of germ cells attracted to the ectopic domain was indeed increased and significantly more germ cells accumulated at the ectopic domain boundary (Fig 6M-Q). Despite the large number of germ cells being attracted we did not observe germ cells or remnants of dying germ cells inside the ectopic domain making it unlikely that germ cells were entering the ectopic domain and subsequently dying. These data show that *wunens* can repel germ cells and prevent them from entering an *hmgcr* expressing ectopic domain.

Taken together we conclude that neither the *wunens* nor the *hmgcr* pathway is dominant and germ cells position themselves using the information provided by both pathways simultaneously.

### *hh* is not required downstream of *hmgcr* for germ cell attraction

We wanted to test if *hh* is downstream of *hmgcr* in attracting germ cells. We therefore asked whether germ cells could be attracted to ectopic *hmgcr* in a *hh* null background. If *hh* is the attractant downstream of *hmgcr* we would predict that germ cells would not be attracted to *hmgcr* in a *hh* background. On the other hand, if *hh* is not the downstream attractant we would predict that germ cells would still be attracted to the *hmgcr* ectopic domain in a *hh* mutant.

We used a null allele, *hh*^*AC*^, that when homozygous causes embryos to have cuticles with a characteristic strong *hh* phenotype consisting of a continuous lawn of denticles identical to that published in Lee et al. 1992 (Fig 7F, G). *hhAC* embryos have very severe patterning defects that are evident from stage 13 which causes germ cells to scatter over the poorly-patterned posterior of the embryo. Therefore we looked earlier in *hhAC* embryos, at stage 12, when germ cells are mostly on track and none have mismigrated into a control ectopic domain that expresses just GFP (Fig 7A, B). In contrast we find that ectopic expression of *hmgcrGFP* is able to attract germ cells in *hh* homozygous mutant embryos similar to sibling heterozygous controls (Fig 7C-E). We conclude therefore that *hh* is not downstream of *hmgcr* in attracting germ cells.

**Figure 7.**
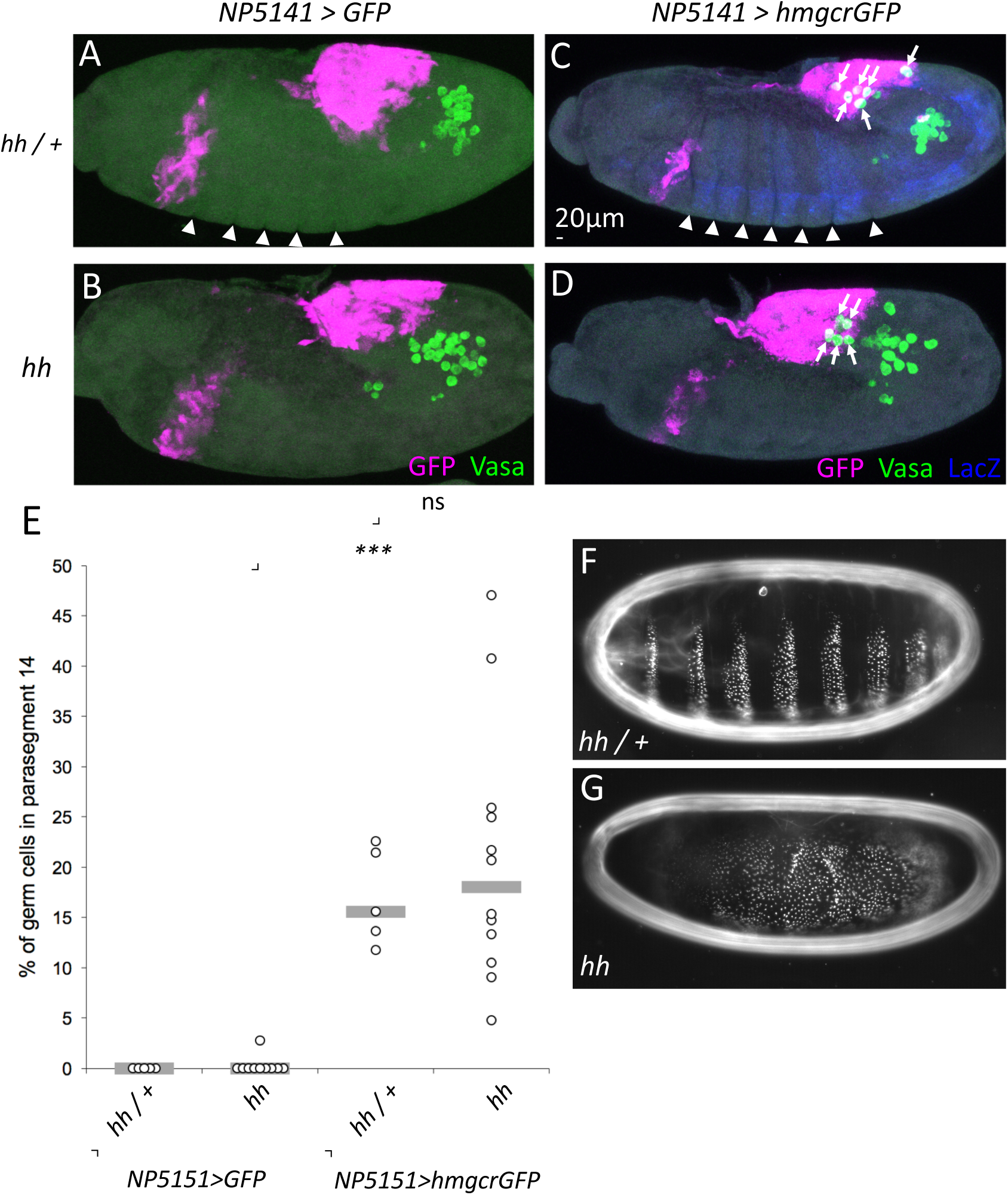
Hh is not downstream of *hmgcr* for germ cell attraction. (A-D) Maximum intensity projections of lateral views of representative stage 12 embryos of the genotypes: (A) *NP5141Gal4/UASGFP; hhAC/+,* (B) *NP5141Gal4/UASGFP; hhAC/hhAC,* (C) *NP5141Gal4/NP5141Gal4; UAShmgcrGFP hhAC/TM3 ftz>lacZ*, (D) *NP5141Gal4/NP5141Gal4; UAShmgcrGFP hhAC*/*UAShmgcrGFP hhAC* fluorescently stained with antibodies against Vasa to label the germ cells (green) and GFP to visualise the ectopic domain (magenta). Heterozygous and homozygous *hhAC* embryos were distinguished by staining with an antibody against LacZ (blue, for C and D) and by loss of wild type parasegmental furrows in the germ band (arrowheads in A and C). Germ cells located in parasegment 14 (the posterior NP5141 expressing domain) are marked with arrows. (E) Graph showing number of germ cells located in parasegment 14 (the posterior NP5141 expressing domain) as a percentage of the total for individual embryos of the genotypes depicted in A-D. Median values are indicated by a grey bar. A Mann-Whitney U test was used to test for statistical significance between the distributions and *p* values are indicated (ns = not significant). (F-G) Darkfield images of cuticle preparations from late embryos from the *UAShmgcrGFP hhAC/TM3 ftz>lacZ* stock showing heterozygous sibling (F) and homozygous *UAShmgcrGFP hhAC* mutant embryos showing a denticle pattern characteristic of *hh* null mutants (G).

## Discussion

Here we have examined the range of influence of a signal downstream of the *hmgcr* pathway that attracts germ cells in *Drosophila* embryos. We have found that this signal can act at distances of at least 51µm. This distance is greater than the distance of virtually all of the germ cells from the target SGPs at stages 10 and 11 and therefore, distance-wise at least, should be sufficient to attract germ cells to the gonad. Furthermore, the signal operates at the same time as a second pathway, namely the Wunens, that influences germ cell migration. This is most strikingly demonstrated by the simultaneous overexpression of both pathways in the same ectopic domain that produces a phenotype different to either pathway alone, in that we see both simultaneous attraction and repulsion. Finally using *hh* null mutants and ectopic *hmgcr* expression we provide evidence that the extracellular signaling molecule Hh is not the chemoattractant downstream of *hmgcr*.

Our 51µm estimate of the range of the *hmgcr*-mediated signal represents approximately 7 germ cell diameters which we would define it as a long-range signal. This range is broadly in line with other long distance signaling molecules both in *Drosophila* and in other species: In *Drosophila* imaginal wing discs the TGF-beta family member Dpp acts at long range, influencing cells up to 20 cell diameters away (Nellen et al., 1996). In Xenopus embryos, TGF-beta ligands can been detected from 7-10 cell diameters away from their source (McDowell et al., 2001) (Williams et al., 2004), whilst in zebrafish embryos cells can be observed responding to endogenous TGF-beta (nodal) signaling at distances up to 200µm (Harvey and Smith, 2009).

Wnt ligands on the other hand can act at either short or long-range. Wingless acts as a short-range inducer in *Drosophila* embryos being secreted by stripes of ectodermal cells and being received only by their neighbours (van den Heuvel et al., 1989). In mouse organoids Wnt3 also acts at short range and could only be visualised only up to 1-2 cells away from synthesizing cells (Farin et al., 2016). On the other hand, Wingless in *Drosophila* imaginal wing discs acts at long range, influencing cells 20 or more cell diameters away (Zecca et al., 1996), and EGL-20 in *C.elegans* can be seen in a gradient up to 50 µm from its source (Coudreuse et al., 2006).

In all of these examples, the TGF-beta or Wnt ligands are providing positional information to static target cells by inducing dose-dependent transcriptional responses. In the case of the Hmgcr and Wunen pathways however the responding cells, namely the germ cells, are motile and a transcriptional response seems unlikely given both the speed at which they respond and the need for the germ cells to response directionally to the signal not just to its strength.

We have demonstrated the distance over which *hmgcr* is potentially able to operate via over-expression of *hmgcr*. The range we have estimated is likely to be influenced by the amount of over-expression of *hmgcr*, in terms of number of cells over-expressing *hmgcr* and by the absolute expression level in each of these cells. For ease of analysis we chose a Gal4 driver that is expressed in the very posterior of the embryo, in parasegment 14, in both the ectoderm and mesoderm. Therefore, the number of cells over-expressing *hmgcr* will be substantial. We estimate there are just over 1000 *hmgcrGFP* ectopically expressing cells in parasegment 14 at stage 10 when using the NP5141 driver (Figure S2A). This would appear to be far greater than the number of cells endogenously expressing *hmgcr* in wild type. For example at stage 12 *hmgcr* is highly expressed in the SGPs (Van Doren et al., 1998) of which there are only 25-35 cells in total per gonad (Sonnenblick, 1941). However earlier, at stages 9-10, *hmgcr* is expressed more broadly in the mesoderm (Van Doren et al., 1998). We estimate there are approximately 250 *hmgcr* expressing mesodermal cells at stage 10 that lie dorsally to the germ cells, and to which the germ cells will migrate (Figure S2B). Therefore, the number of cells ectopically versus endogenously expressing *hmgcr* is comparable at early stages at least.

There are two possible models of the interactions between Hmgcr and Wunen (Fig 8). The prevailing view has been a two signal model (Fig 8A) (Barton et al., 2016). One chemoattractant results from Hmgcr expression in the mesoderm and is perceived by germ cells via an unidentified receptor. The second chemoattractant is perceived by germ cells using the Tre1 GPCR. It is also a substrate for the Wunens and is dephosphorylated and thereby destroyed by Wunen expressing cells, including the germ cells, which collectively act as a sink for the chemoattractant. In this model, the spatial information provided by Hmgcr and Wunens is integrated at the level of the germ cells which use the information provided by both chemoattractants.

**Figure 8.**
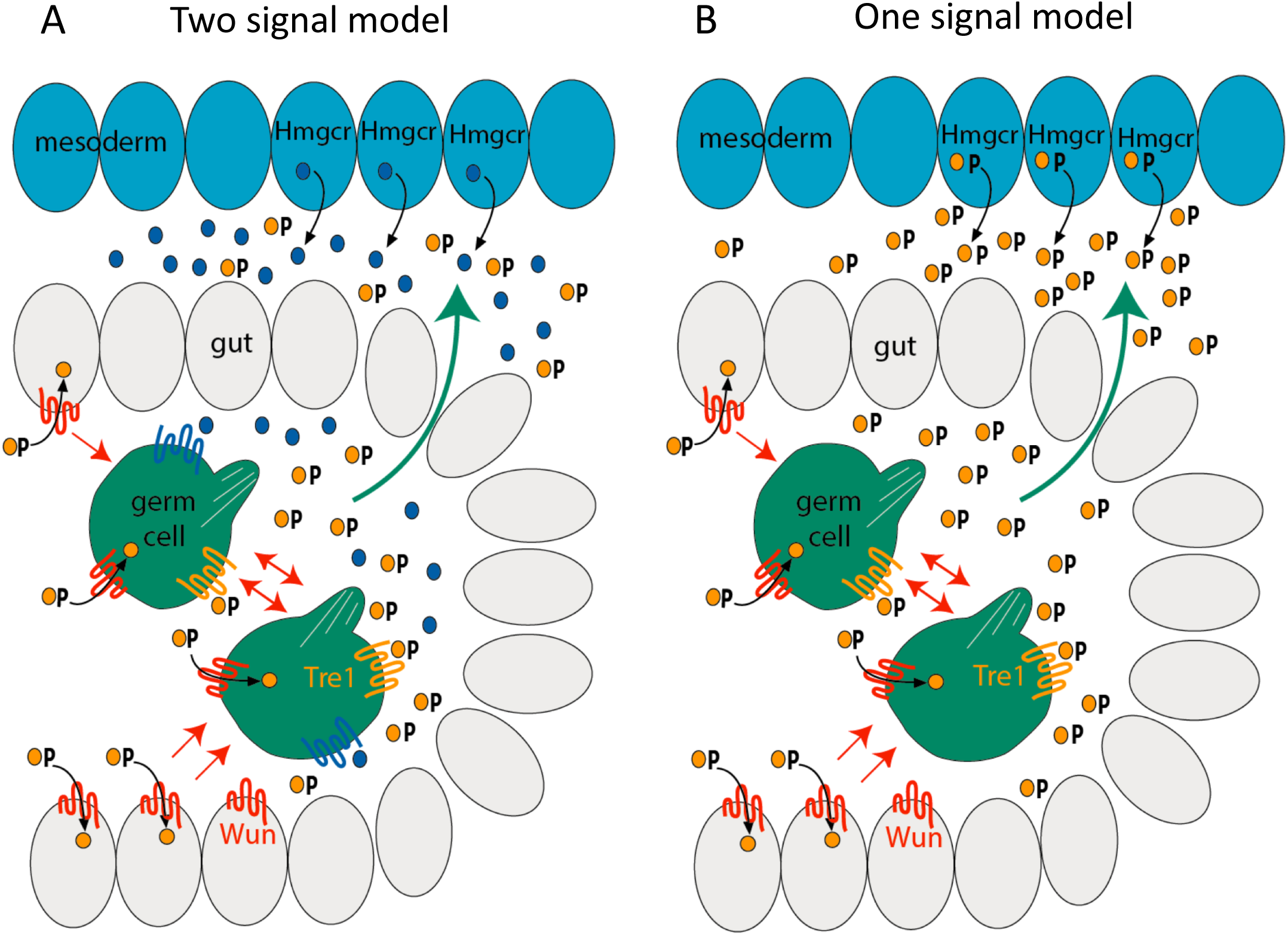
One and two signal models of how Wunen and Hmgcr mediate germ cell migration. Schematic of lateral views of a stage 9 embryo with germ cells inside the posterior midgut being attracted (green arrow) towards Hmgcr expressing mesodermal cells (blue cells) and being repelled from regions of somatic Wun expression and each other due to germ cell Wun expression (red arrows). (A) In a two signal model (Barton et al., 2016), Hmgcr expression results in release of a chemoattractant (blue circle), unrelated to Wun, which is detected on germ cells via a receptor other than Tre1 GPCR (blue receptor). Localised Wunen expression (on germ cells and some somatic cells) acts as a sink for a second chemoattractant containing a phosphate group (yellow circle) via dephosphorylation. This second chemoattractant is detected on germ cells by the Tre1 GPCR (yellow receptor). (B) In a one signal model, Hmgcr expression results in release of a chemoattractant containing a phosphate which is detected on germ cells via Tre1 and dephosphorylated by Wunens which again act as a sink.

In a single signal model Hmgcr expression results in secretion of a chemoattractant from the mesoderm which is also the substrate for the Wunens and is detected on germ cells by the Tre1 GPCR (Fig 8B). In this model, the spatial information provided by Hmgcr and Wunens is integrated at the level of the chemoattractant gradient which depends on the combined actions of both of these enzymes.

Both models have precedents from other extracellular gradients both in *Drosophila* and other organisms. The one signal model (Fig 8B) resembles classical source-sink models for both chemoattractant and morphogen gradients (Cai and Montell, 2014). The use of simultaneous attraction and repulsion, as per the two signal model (Fig 8A), is seen in *Drosophila* axonal pathfinding where commissural axons are attracted and repelled by the ligands Netrin and Slit respectively (reviewed in Dickson and Gilestro, 2006). The migration of vertebrate trunk neural crest cells is controlled by both positive and negative regulators including ligand/receptor pairs such as ephrin/Eph, and Sdf1/Cxcr4 (reviewed in Shellard and Mayor, 2016). Both of these examples differ slightly to our model because repulsion is occurring via direct detection of a chemorepellent rather than via destruction of an attractant.

Based on published observations and the data presented here we now favour the one signal model (Fig 8B) for a number of reasons. Firstly, we have shown that Hh is not downstream of the *hmgcr* pathway for attracting germ cells. If this had been the case it would have made the two signal model highly likely, because Hh has not been reported to be phosphorylated (Lee et al., 2016) and therefore not likely to be a substrate of the Wunens. Secondly, the signals downstream of both the Wunens and Hmgcr operate over similar long-ranges which means they are potentially the same. Thirdly, zygotic loss of function mutants of *wun* and *clb* both exhibit similar very strong mis-migration phenotypes (Van Doren et al., 1998; Zhang et al., 1997). If each pathway influenced their own independent signal, then one might expect that removal of either pathway alone would result in partial germ cell mis-migration as the other would still be active and also acts over a long range. Fourthly, we have examined whether the *wunen* and *hmgcr* pathways act simultaneously rather than consecutively. If the pathways had acted consecutively, this would only be compatible with the two-signal model. We found instead that the pathways act simultaneously (Fig 6M-P), a result consistent with the one signal model, but does not exclude the two signal model. Finally, we showed that the *hmgcr* and *wunen* pathways actually do not operate in isolation, as per the two signal model, because we see decreased germ cell attraction to ectopic *hmgcr* in a *wun* mutant background (Fig 2), therefore favouring the one signal model.

The ultimate confirmation of which model is correct will require identification of the chemoattractant(s). It is interesting to note that germ cell migration in other species such as chicken and zebrafish seems to require only a single chemoattractant, SDF-1, in spite of the much longer migratory journeys, both in terms of distance and time, in these species (Barton et al., 2016). It is clear that *Drosophila* germ cells can’t be responding to SDF-1 as no SDF-1 homolog exists in flies. What is less clear is whether the signals downstream of *wunens* and *hmgcr* exist in vertebrates, perhaps playing a more subtle role. Tantalising evidence of from zebrafish suggests this might be the case, with simultaneous knockdown of all the Wunen homologues causing some germ cells to mis-migrate (Paksa et al., 2016). Therefore, the cues that regulate *Drosophila* germ cell migration might actually be more conserved than we first thought.

## Author contributions

K.K., A.M. and A.D.R. designed the experiments, K.K. made the *Drosophila* stocks and K.K. performed the experiments with the exception of the *hmgcr in situ* hybridisations which were performed by A.M. A.D.R. did the tracking for the live imaging, A.D.R. wrote the paper.

## Acknowledgements

We thank Tom Starkey for producing some of the recombinant fly stocks, the University of Nottingham School of Life Sciences Microscope facility (SLIM) for use, maintenance and training on the Zeiss 880 confocal microscope and Malcolm Bennett and Antony Bishopp (School of Biosciences, University of Nottingham) for use and training on the Zeiss Z1 light sheet microscope (funded through BBSRC award BB/M012212/1). We also thank Christian Feldhaus (Max Planck Institute for Developmental Biology) for assistance with distance measurements with Imaris and Markus Owen (School of Mathematics, University of Nottingham) and Andrew Johnson and Fred Sablitzky (School of Life Sciences, University of Nottingham) for comments on the manuscript. We also acknowledge the Flybase consortium for information on genes and fly stocks and the Bloomington stock centre at Indiana University for fly stocks. We thank the lab of Saverio Brogna for reagents and the University of Nottingham for the studentship for KK. The authors declare no conflicts of interest.

## Supplementary Materials

**Figure S1.**
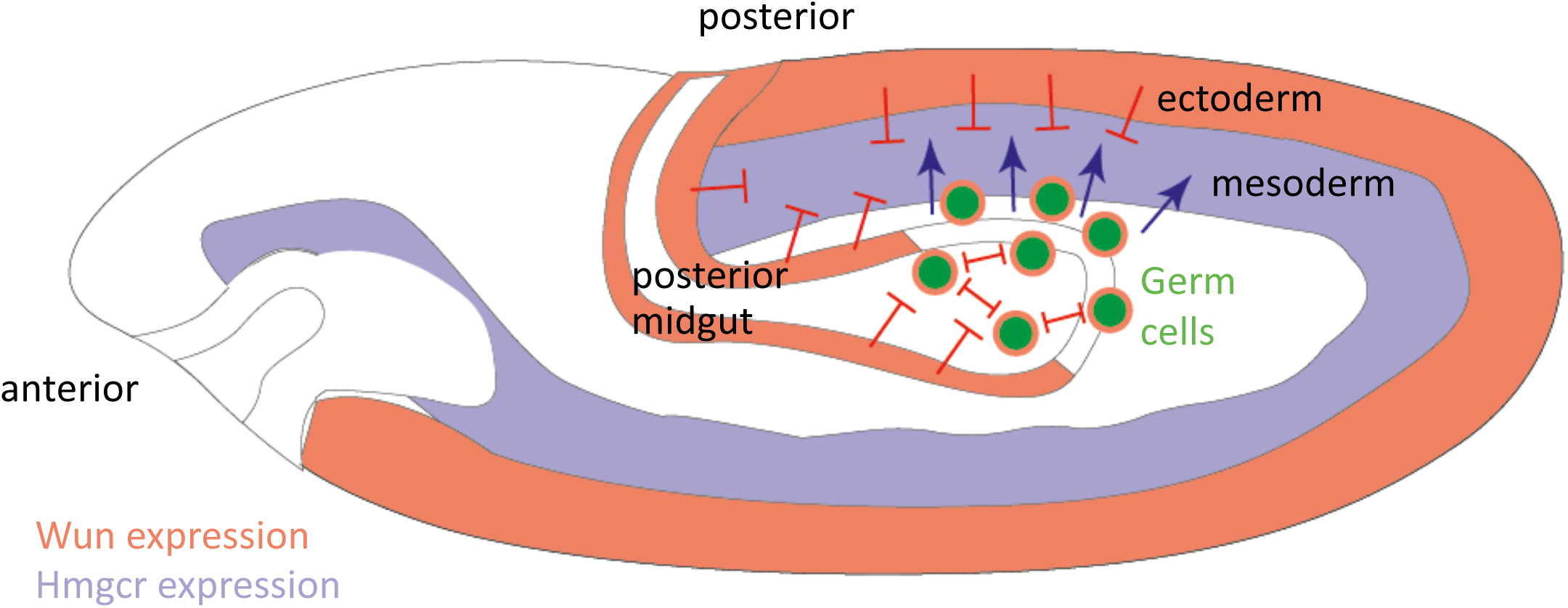
Complementary expression patterns of Hmgcr and Wun guide germ cells to the SGPs. Hmgcr expression (light blue) in the mesoderm and later just in the SGPs, which are derived from the mesoderm, is responsible for attracting (blue arrows) the germ cells (green). Wunen expression (light red) in regions of the ectoderm and posterior midgut repels germ cells (red bar-headed lines). Wunen expression on germ cells themselves causes mutual repulsion leading to germ cell dispersal.

**Figure S2.**
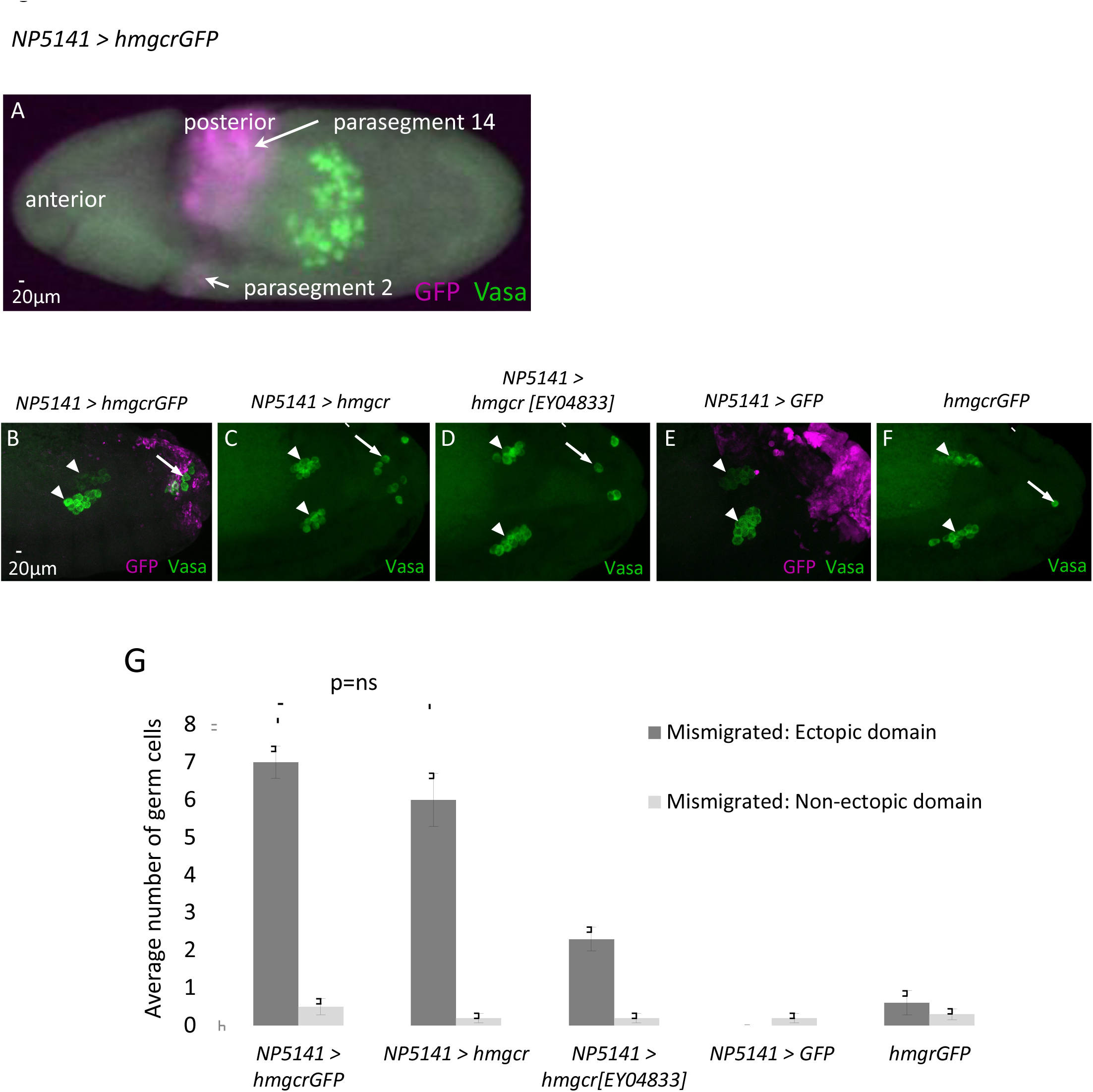
GFP tagged Hmgcr is functional. (A) Maximum intensity projection of a lateral view of a *NP5141Gal4/+; UAS hmgcrGFP/+* stage 10 embryo, fluorescently stained with antibodies against Vasa to label the germ cells (green) and GFP to visualize the *hmgcrGFP* expression (magenta). *NP5141*-driven transgene expression is observed in parasegments 2 and 14. (B-F) Maximum intensity projections of lateral views of stage 14 embryos, fluorescently stained with antibodies against Vasa (green, germ cells) and GFP (magenta, ectopic domain) (B, E only) of the genotypes: *NP5141Gal4/+; UAShmgcrGFP/+* (B), *NP5141Gal4/+; UAShmgcr/+* (C), *NP5141Gal4/+; hmgcrEY04833/+* (D), *NP5141Gal4/UASGFP* (E) and *UAShmgcrGFP/UAShmgcrGFP* (F). Arrows indicate mismigrated germ cells located in the ectopic domain, arrowheads indicate the embryonic gonads. Estimate of boundary of ectopic domain indicated with dashed white line (C, D, and F). (G) Graph showing the mean ± s.e.m number of mismigrated germ cells (defined as being outside of the gonad cluster) located either within or outside of the ectopic domain in stage 14 embryos. n=10 embryos scored per genotype. P value calculated by Student’s t-test.

**Figure S3.**
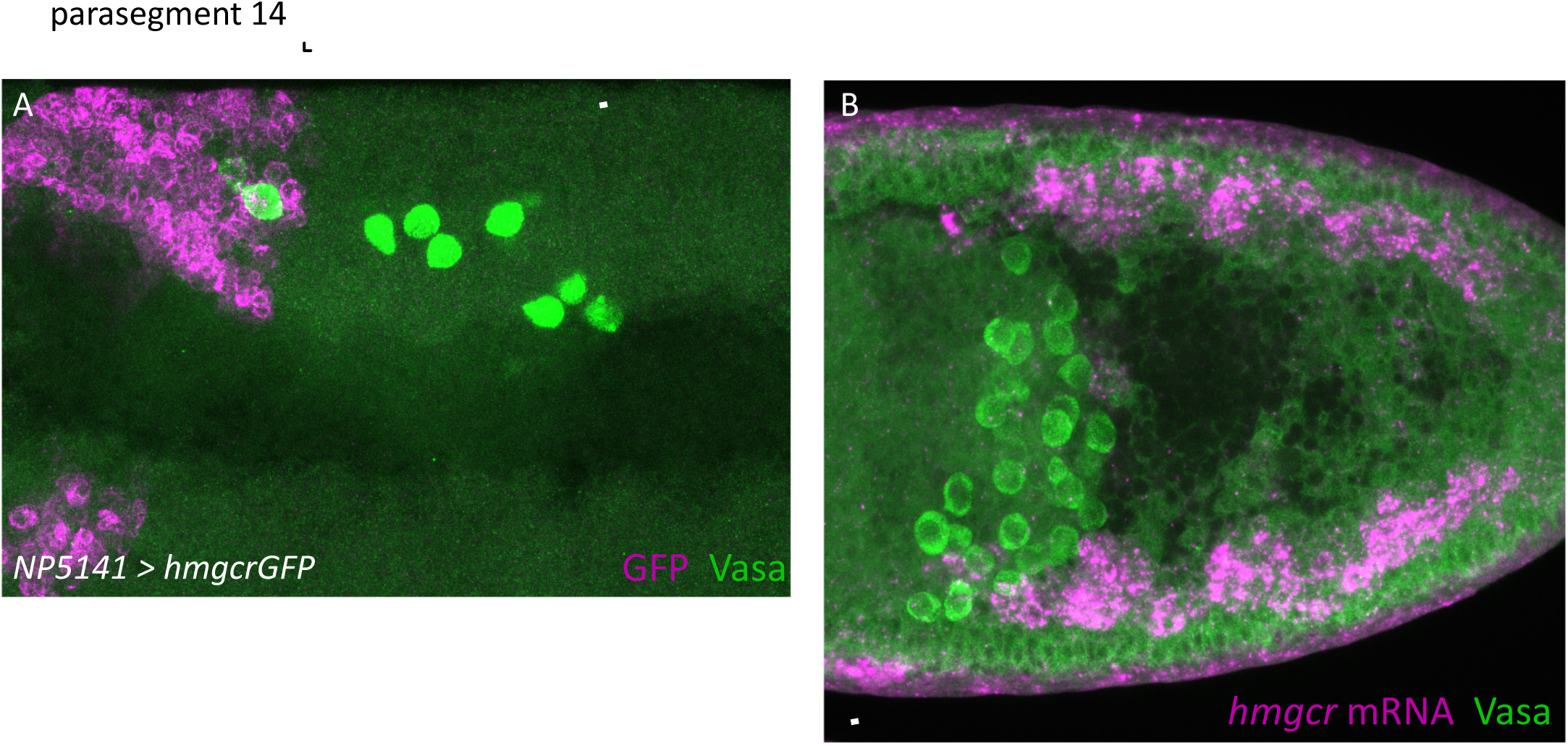
Images used to estimate the number of *hmgcr* endogenously and ectopically expressing cells. (A) Lateral view of a maximum intensity projection of a *NP5141Gal4/+; UAS hmgcrGFP/+* stage 10 embryo, fluorescently stained with antibodies against Vasa to label the germ cells (green) and GFP to visualise the ectopic domain (magenta). The number of *hmgcr* expressing cells in parasegment 14 was scored manually. (B) Maximum intensity projection of a dorsal view of a wild type embryo at stage 10 fluorescently stained with antibodies against Vasa to label the germ cells (green) and a *hmgcr* RNA probe to visualise *hmgcr* expression (magenta). The number of *hmgcr* expressing cells in the mesoderm that overlies the germ cells was scored manually. Scale bars are 50 µm.

